# RNA i-Motif Formation at Neutral pH

**DOI:** 10.64898/2026.03.01.708865

**Authors:** Zara Preciado, Sophie Fountain, Elisé P. Wright, Dilek Guneri, Mahmoud A. S. Abdelhamid, Timothy Craggs, Yiliang Ding, Zoë A. E. Waller

**Affiliations:** School of Chemistry, Pharmacy and Pharmacology, UEA, Norwich Research Park, NR4 7TJ, UK; The University of Sheffield, Department of Chemistry, Western Bank, Sheffield, UK; School of Pharmacy, University College London, 29-39 Brunswick Square, London, WC1N 1AX, UK; School of Health, Science and Society, University of Suffolk, 19 Neptune Quay, Ipswich, IP4 1QJ, UK; Department of Cell and Developmental Biology, John Innes Centre, Norwich Research Park, Norwich NR4 7UH, UK

## Abstract

i-Motifs are non-canonical nucleic acid secondary structures formed in sequences rich in cytosine. Previous work has shown that DNA i-motifs may form at neutral pH and provided evidence to suggest they influence biological functions. RNA i-motifs are less stable than DNA i-motifs, so more questions surround the biological relevance of these structures in RNA. Using biophysical methods, we found that increasing cytosine tract lengths resulted in increased thermal stability but not pH stability. In ensemble solution experiments RNA i-motifs appear globally unfolded at neutral pH. However, single molecule experiments revealed that 1% of RNA i-motifs remain folded in solution. This has implications for the potential of formation of RNA i-motifs in cells.

## Introduction

i-Motifs are non-canonical nucleic acid structures formed from cytosine rich sequences stabilised by hemi-protonated cytosine-cytosine base pairs.^1^ i-Motifs are relatively unique nucleic acids structures, due to the requirement of hemi-protonation of the cytosines for stable formation.^2^ It has been shown that targeting i-motif DNA structures in gene promoters can modulate gene transcription.^3, 4^ Our own work showed that certain DNA sequences may fold into i-motifs at neutral pH, and examples of such sequences are present throughout the human genome.^5^ Evidence using in cell NMR and specific antibodies have also demonstrated the presence of i-motifs in cells.^6-8^

Early work into the structure of i-motifs determined that although formation of i-motifs in DNA was possible, even if at a low proportion of folded species,^9^ formation of RNA i-motif structures at neutral pH is not possible.^10^ RNA i-motif structures can fold at acidic pH, but they are globally unfolded at pH 7. The reasons for this is fundamentally down to the sizes of the grooves. i-Motif structures have two wide grooves and two narrow grooves. The wide grooves are wider than B-form DNA, providing a large surface different to that of other DNA structures. The minor grooves in i-motif are very narrow (3.1 Å), approximately half the distance of B-DNA.^11^ In the narrow grooves the position of the sugar-phosphate backbones are such that in DNA there are phosphate-phosphate interactions.^4^ The additional 2’-OH group in the equivalent RNA structures provides both steric and electronic clashes, in line with their overall stability compared to DNA i-motifs. Consequently, there is repulsion between adjacent negatively charged phosphate backbones. The 2’-OH groups in RNA i-motifs destabilise i-motifs because of the close juxtaposition of sugar phosphates in this structure, making formation of RNA i-motif disfavoured. It has been proposed that sugar–sugar contacts are important in stabilising i-motifs in general and reveal that solvation is the cause of the instability of RNA i-motif structures.^12^ This general difference in stability between DNA and RNA i-motif structures is in direct contrast to G-quadruplexes, where RNA G-quadruplexes are typically much more stable than their DNA counterparts.^13^

Despite the instability of RNA i-motifs *in vitro*, experiments by Christ and co-workers using the i-motif specific antibody iMab showed that these structures may exist in RNA in cells.^8^ They showed the presence of foci in the cytoplasm, indicating potential formation of RNA i-motifs, that are not removed by DNase I digestion. A small decrease in the number of iMab foci after RNase A treatment supports this speculation.

In our work on DNA i-motifs we determined that longer C-tracts (and stacks of CC base pairs) gave rise to more stable i-motifs.^5^ We also determined a point where sequences became stable at neutral pH. Given the increasing interest in i-motifs in general, the lack of general characterisation of RNA i-motifs and the evidence from Christ’s group that showed that RNA i-motifs may be present in cells,^8^ we questioned whether RNA i-motifs could form at all *in vitro*. Herein we show that RNA i-motifs can form at neutral pH and are present at proportions of 1% in solution using single molecule FRET.

## Materials and Methods

### RNA

RNA sequences were supplied by Eurogentec (Belgium), synthesized on a 1000 nmol scale and purified by reverse phase HPLC. Samples were dissolved in ultrapure water to give approximately 1 mM final concentrations, confirmed using a Nanodrop and diluted to 1 mM as appropriate. Each RNA sequence in the library consisted of four cytosine tracts separated by three loops containing uracils. The notation C_n_U_x_ is used where *n* = the number of cytosines in each of four tracts and x = the number of uracil in each of the three loops. A summary of the model RNA sequences used in this study is given in Table 1. For all experiments, RNA sequences were diluted in buffer containing 10 mM sodium cacodylate and 100 mM sodium chloride at the specified pHs. Annealing was performed by heating in a heat block at 95°C for 5 min and samples were cooled slowly to room temperature overnight for use the following day.

**Table 1.** Library of oligonucleotides sequences and data showing melting (*T*_m_) and annealing (*T*_a_) temperatures for the sequence [C_n_U_3_]_3_C_n_ measured in 10 mM sodium cacodylate with 100 mM sodium chloride including the average for the melting and annealing temperatures and standard deviation for 3 cycles. *nd: not discernible.

| Sequence | Bases | Notion | $T_m$ at pH 5.5 | $T_a$ at pH 5.5 | pH<sub>T</sub> |
| --- | --- | --- | --- | --- | --- |
| $[C_1U_3]_3C_1$ | 13 | C1U3 | nd | nd | nd |
| $[C_2U_3]_3C_2$ | 17 | C2U3 | $7.3 \pm 0.6$ | $8.0 \pm 0.9$ | 5.0 |
| $[C_3U_3]_3C_3$ | 21 | C3U3 | $16.2 \pm 0.6$ | $14.1 \pm 1.7$ | 5.5 |
| $[C_4U_3]_3C_4$ | 25 | C4U3 | $18.8 \pm 0.9$ | $17.4 \pm 0.6$ | 5.4 |
| $[C_5U_3]_3C_5$ | 29 | C5U3 | $21 \pm 0.9$ | $19.4 \pm 0.6$ | 5.5 |
| $[C_6U_3]_3C_6$ | 33 | C6U3 | $25.1 \pm 0.4$ | $21.3 \pm 0.6$ | 5.6 |
| $[C_7U_3]_3C_7$ | 37 | C7U3 | $27.7 \pm 0.8$ | $21.6 \pm 0.6$ | 5.6 |
| $[C_8U_3]_3C_8$ | 41 | C8U3 | $29.7 \pm 0.7$ | $21.3 \pm 0.6$ | 5.6 |
| $[C_9U_3]_3C_9$ | 45 | C9U3 | $32.2 \pm 0.7$ | $22.6 \pm 0.6$ | 5.6 |
| $[C_{10}U_3]_3C_{10}$ | 49 | C10U3 | $37.0 \pm 0.6$ | $24.2 \pm 0.3$ | 5.6 |

### UV absorption spectroscopy

UV spectroscopy experiments were performed on a Cary 60 UV-Vis spectrometer (Agilent Technologies) equipped with a TC1 Temperature Controller (Quantum Northwest) and recorded using a low volume masked quartz cuvette (1 cm path length). ODNs (Tables 1) were diluted to 2.5 μM in buffer at the desired pH. Samples (200 μl) were transferred to a cuvette, covered with a layer of silicone oil and stoppered to reduce evaporation of the sample. The absorbance of the RNA sequence was measured at 295 nm as the temperature of the sample was held for 10 min at 4°C then heated to 95°C at a rate of 0.5°C per min, held at 95°C before the process was reversed and the samples cooled back to 4°C. Each melting/annealing process was repeated three times. Data were recorded every 1°C during both melting and annealing and each point was the average of three scans. Melting temperatures (*T*_m_) were determined using the first derivative method. Thermal difference spectra (TDS) were calculated by subtracting the spectrum between 220 and 320 nm of the folded structure at 4°C from that of the unfolded structure at 95°C. The data was normalized and the maximum change in absorption was set to +1 as previously described (<u>27</u>).

### Circular dichroism

CD spectra were recorded on a Jasco J-810 spectropolarimeter using a 1 mm path length quartz cuvette. RNA sequences (Table 1) were diluted to 10 μM (total volume: 200 μl) in buffer at pH increments of 0.5 pH unit from 4.0 to 8.0. The scans were recorded at room temperature (20°C) between 200 and 320 nm. Data pitch was set to 0.5 nm and measurements were taken at a scanning speed of 200 nm/min, response time of 1 s, bandwidth of 2 nm and the 100 mdeg sensitivity; each spectrum was the average of three scans. Samples containing only buffer were also scanned according to these parameters to allow for subtraction of residual buffer signals. The transitional pH of the i-motif was primarily determined by monitoring the CD signal at 288 nm. In addition, CD data at 200 nm and the maximal ellipticity values were analysed to provide a more comprehensive assessment of the pH-dependent folding behaviour. The transitional pH was calculated from the inflection point of the fitted ellipticity curve at 288 nm. Final data analysis and figure preparation were performed using Origin.

### Single Molecule FRET Experiments

#### Sample Preparation

Labelled RNA was supplied by Eurogentec (Belgium), synthesised on a 200 nmol scale and purified by reverse-phase HPLC. Oligonucleotides were dissolved at approximately 100 μM in MilliQ water and quantified using a Nanodrop. Oligos were diluted 500 nM in 10 mM Sodium Cacodylate (pH 5.5 and 7), 100 mM NaCl. All samples were annealed by heating at 95°C for 5 min and then cooled slowly to room temperature overnight.

#### smFRET Measurements

smFRET measurements were performed using a home-built confocal microscope (smfBox) with alternating laser excitation (ALEX).^14^ Two diode lasers (515 nm and 635 nm, Omicron LuxX Plus) were directly modulated with a 100 µs cycle (45 µs donor excitation, 5 µs off, 45 µs acceptor excitation, 5 µs off) and combined into a single-mode fibre. The output beam was collimated and cropped to 5 mm using an iris before being directed into an Olympus UPLSAPO 60× NA 1.35 oil-immersion objective via a Chroma ZT532/640rpc excitation dichroic. Fluorescence emission was collected by the same objective, focused through a 20 µm pinhole, spectrally split using a 640 nm long-pass dichroic, and detected by two avalanche photodiodes (Excelitas SPCM-AQRH-14 and SPCM-NIR-14). Photon arrival times were time-stamped using a National Instruments PCIe-6353 card, with acquisition controlled by EI-FLEX software (REF).^15, 16^Annealed samples were diluted to ∼5 pM in observation buffer (10 mM Sodium Cacodylate (pH 5.5 and 7), 100 mM NaCl, 100 mM NaCl; 0.1 mg/mL photobleached BSA), and measurements were acquired for 60 minutes in triplicate. Extended measurements were then conducted at both pH values for 3 hours. For pH 7, an additional 12-hour experiment was performed.

#### Data Analysis

Data analysis was performed with Jupyter Notebooks using the FRETBursts Python module (version 0.7.1).^17^ Background in each channel was estimated using an exponential fit of inter-photon delays. Bursts were identified by performing a dual-channel burst search (DCBS) using a photon sliding window algorithm with m = 10, F = 20 (for each channel) and DD+DA and AA thresholds = 30, to extract doubly labelled bursts from each acquisition.

#### Burst Variance Analysis (BVA) for FRET Distributions

Burst Variance Analysis (BVA) was performed using FRETBursts to compute the standard deviation of sub-burst FRET efficiencies for each photon burst, allowing identification of static versus dynamic FRET populations by comparing observed fluctuations to those expected from the shot-noise limit. BVA looks for deviations from an expected sub-burst E variance given the E of the whole burst.^17, 18^ On the BVA plot, the standard deviation values clustering around the expected standard deviation indicate static FRET, whereas bursts with standard deviation values significantly above the confidence interval indicate within-burst dynamics.

## Results and Discussion

In line with previous work on DNA i-motifs,^5^ we decided to vary the length of the C-tracts in sequences, whilst keeping the loop length constant to determine whether there was a sequence which was stable at neutral pH. The sequences examined are shown in Table 1.

UV spectroscopy was used to determine both the melting (*T*_m_) and annealing temperatures (*T*_a_) for each of the RNA sequences (Figure S1, S2). All of the sequences in our model library were thermally annealed and melted in 10 mM sodium cacodylate with 100 mM sodium chloride at both pH 5.5 and 7.4. These pH conditions were selected to offer an acidic environment for i-motif formation (5.5) and physiological pH for comparison (7.4). Using these conditions allowed us to directly compare with our previous work on the equivalent DNA sequences.^5^

At pH 5.5, all the sequences from the RNA library showed melting and annealing curves when monitored at 295 nm except C1 (Figure S1). This is indicative that the sequences form i-motif structures, as this wavelength is characteristic of i-motif formation. As the number of cytosines in each tract increases, the stability of the structures formed also increases (Figure S1 and S2). This indicates that the additional cytosines in the stack of CC base pairs offers some increase in overall stability.

We used thermal difference spectroscopy to determine the thermal difference spectra of the sequences examined, which indicated that at pH 5.5 all but C1 formed spectra with a positive peak at 240 nm and a negative peak at 295 nm, indicating i-motif structure formation (Figures S3 and S4).^19^

Analogous experiments repeated at pH 7.4 revealed the UV melting/annealing profiles did not show any transition or melting points (Figure S12). This is in contrast to DNA, where i-motif structures can form in sequences with 5 or more cytosines tracts at pH 7.4.^5^ This is indicative that no sequence examined gave predominant signals indicative of formation of i-motif structures at physiological pH.

To complement the TDS, we measured circular dichroism spectra for the sequences at different pHs and determined the pH stability by measurement of the transitional pH of i-motif formation (pH_T_, Figures S5-11). To do this, each sequence was measured in 10 mM sodium cacodylate with 100 mM sodium chloride at a range of pHs between 4.0 and 7.0. Example data is given in Figure 1, calculated transitional pHs are provided in Table 1.

**Figure 1:**
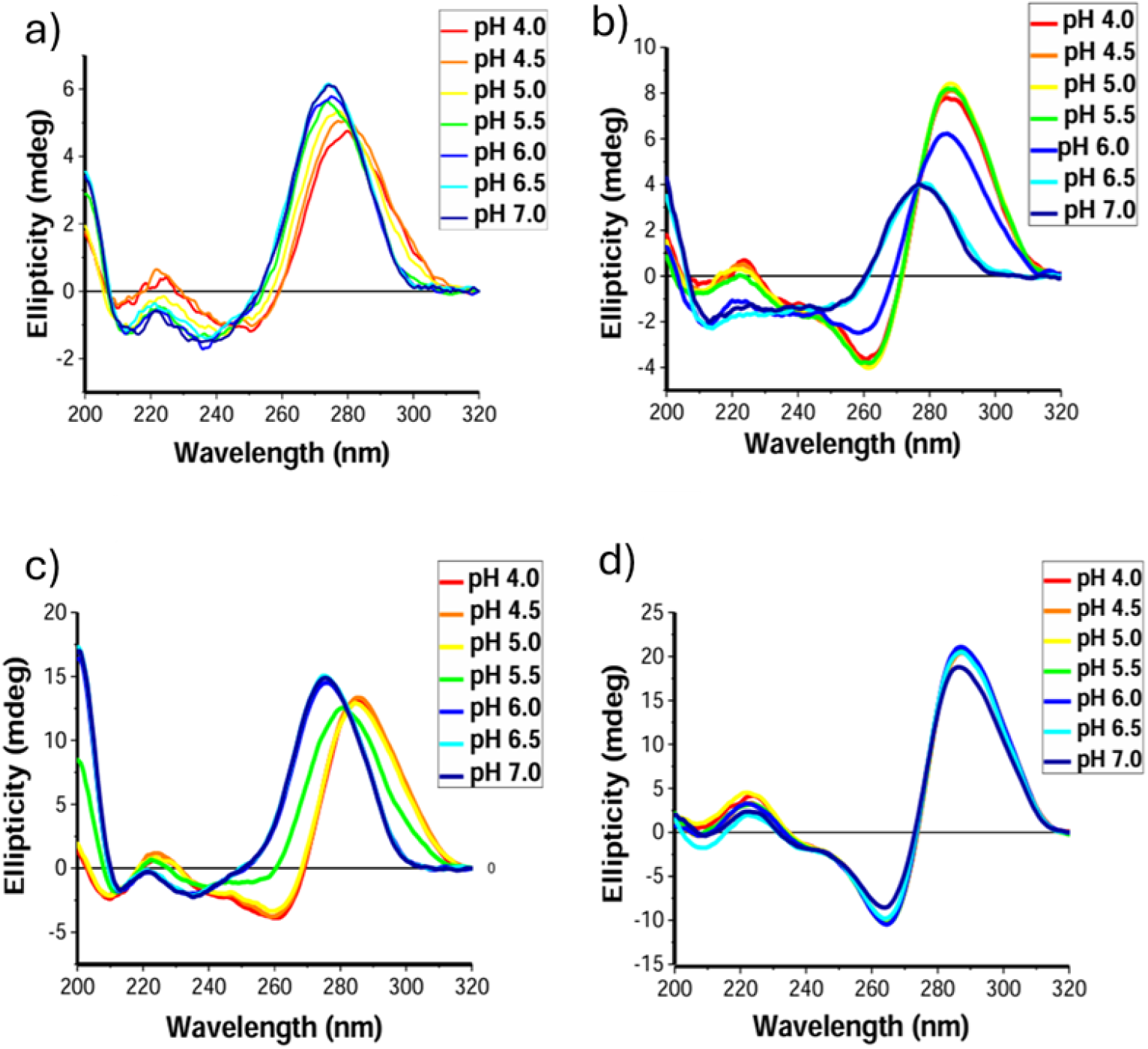
Circular Dichroism of 10 µM sequence in 10 mM sodium cacodylate buffer, 100 mM NaCl at the specified pHs (a)C2U3(RNA), (b)C2T3(DNA), (c)C5U3(RNA), (d)C5T3(DNA)

From the CD of the RNA sequences it is possible to see differences in the CD spectra when compared to the DNA analogues. In i-motif forming DNA sequences at acidic pH, there is a positive peak at 288 nm (Figure 1b/d), which then shifts to 275 nm, and decreases in intensity as the pH increases as the i-motif unfolds.^20^ In the analogous RNA sequence, there is still a positive band at 288 nm at acidic pH, indicative of formation of i-motif structures but when the pH increases, and the i-motif unfolds, rather than a shift towards lower wavelengths and a decrease in ellipticity, there is a shift, but the ellipticity does not decrease in the same way. This is consistent across all the sequences (Figure SX), indicative that the unfolded C-rich sequences in RNA have different spectroscopic properties than the equivalent sequences in DNA. The transitional pHs were determined by plotting the ellipticity at 288 nm against the pH in which the sequence was measured. From these it is possible to determine that for [C_2_U_3_]_3_C_2_ the transitional pH was determined to be pH 5.0. For every other sequence examined thereafter the transitional pH was measured to be the approximately the same: pH 5.5. There was no observed trend or change that occurred when increasing the number of cytosines in the tract/stack. This is in direct contrast with the equivalent sequences in DNA, which showed an increase in pH stability with increasing number of cytosines in the tract lengths.^5^ For the sequences observed, this indicated that the pH stability is capped at pH 5.5, regardless of sequence. Knowing the structural constraints in the narrow grooves of i-motif structures, this is unsurprising. It leaves us the question as to the results observed by Christ’s group, where they used the i-motif specific antibody to show that i-motifs were observed in the cytoplasm of cells.^8^ These were not removed by DNase I digestion and a small decrease in the number of iMab foci after RNase A treatment supported the speculation that these were RNA i-motifs. We considered that there may indeed be endogenous compounds or conditions in cells that may further stabilise RNA i-motifs. We also questioned whether there were potentially a small amount of folded RNA i-motifs in solution at neutral pH which were masked by the ensemble CD experiments. To test this, we examined an exemplar RNA sequence C5U3_FRET_ (donor) 5’-Atto550-CCCCCUUUCCCCCUUUCCCCCUUUCCCCC-3’-Atto647N (acceptor) using single molecule FRET experiments (Figures S14-18) There were some challenges with this sequence, because the dyes are at the ends of the C-rich tracts, without any flanking sequences. This does result in a significant amount of quenching (Figure S14B) because the fluorophores overlap when the i-motif is formed. This also means that it is more difficult to detect folded structures. Burst variance analysis was performed to determine whether the heterogeneity at the different FRET efficiencies is due to dynamics. Burst variance analysis (BVA) is a method that detects within-burst dynamics by comparing the standard deviation of the FRET efficiency of bursts to what is expected. This test distinguishes between static and dynamic sources of heterogeneity in single-molecule FRET histograms by comparing the standard deviation of each molecule observed to the standard deviation expected for a static species at different FRET efficiencies. In this case, when the standard deviation is greater than expected, it’s likely because the molecule giving the FRET efficiency is dynamic. We measured C5U3_FRET_ at both pH 5.5 and pH 7.0 for one hour. At pH 5.5, there is a population of high FRET efficiency (near to 1), and a smaller population at lower FRET efficiency (0.4), consistent with a folded structure where the two ends have come into close proximity (Figure 2A/B). The decrease in the number of detected bursts suggests that the fluorophores may be in very close proximity, leading to dye-dye quenching, which is known to reduce the observable FRET signal.^21^ Even with the quenching issue at pH 7, it is possible to detect a small population of species with high FRET efficiency, indicating that at least 1% of the molecules detected in solution are folded. This indicates that the sensitivity of single molecule methods enables the small, proportion of RNA i-motif which remains folded at neutral pH to be detected even in dilute conditions *in vitro*. We examined the same sample over time, and across 1, 3 and 12 hours the percentage of folded species were 1.58, 1.63 and 1.16% respectively. This indicated that the equilibrium between folded and unfolded i-motif formed by this structure at pH 7 is consistent over time with a stable and detectable proportion of folded structure (Figures S16E-18E).

**Figure 2:**
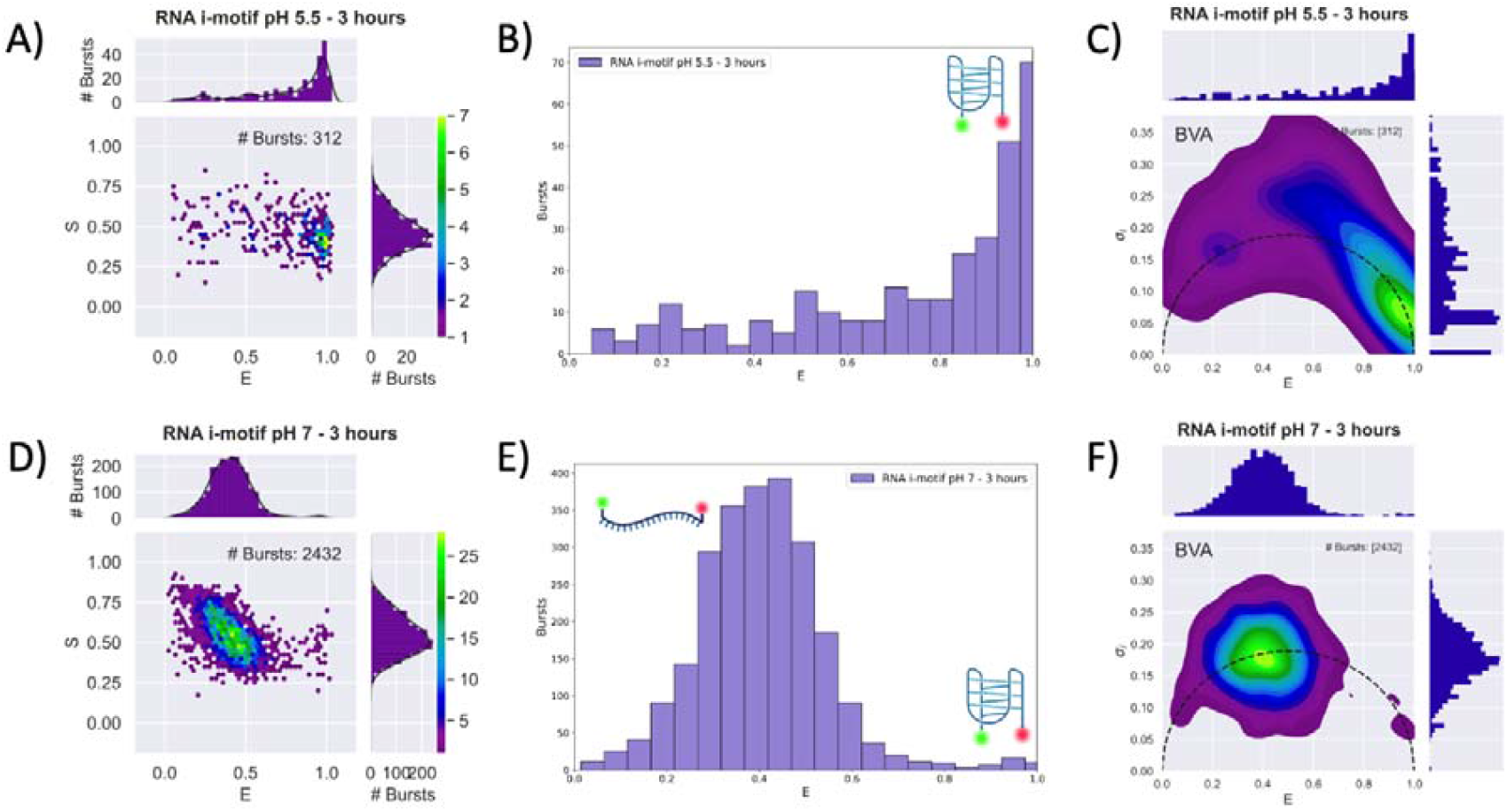
Single Molecule FRET Experiments A) C5U3_FRET_ at pH 5.5 2D-histogram of dual channel burst search B) C5U3 at pH 5.5 histogram of FRET Efficiency from the dual channel burst search C) BVA 2D histogram for C5U3 at pH 5.5 D) C5U3 at pH 7.0 dual channel burst search filtered for doubly labelled molecules E) C5U3 at pH 7.0 histogram of FRET Efficiency from the dual channel burst search F) BVA 2D histogram for C5U3 at pH 7.

For many years researchers avoided working on i-motifs because it was presumed their sensitivity to pH meant that they were not physiologically relevant.^10^ In direct contrast to G-quadruplexes, where RNA G-quadruplexes are more stable than their DNA analogues, RNA i-motifs are significantly less stable than their DNA counterparts.^13^ *T*his does not mean they do not form, but they are only a small proportion of folded species at neutral pH. Given that localised pH varies across the cell, and intracellular pH varies between cell types and age. We propose that RNA i-motifs may play physiological roles in cells and are worthy of further investigation. *T*he wave of work on liquid-liquid phase separation^22, 23^ and interest in membrane-less compartments in cells, coupled with evidence that organelles have pH gradients of their own,^24^ questions the dogma that cells have a fixed intracellular pH.^25^ As the architecture of the insides of cells are revealed in more detail it becomes intuitive that this provides the opportunity for distinct, localised pHs in cells. *T*his is highlighted in recent work from the Krishnan group which demonstrated clearly that intracellular pH gradients exist within both organelles and in membrane-less organelles, even in the absence of ion-transporting proteins.^24^ Moreover the recent disclosure that acidosis orchestrates adaptations of energy metabolism in tumours indicates how changes in pH trigger biological responses.^26^ *T*his opens up the opportunity and scope for RNA i-motif and i-motif-like structures and their potential use as conformational switches in cells.

## Conclusions

Herein we show that RNA i-motifs can form even at near-neutral pH, albeit in small proportions. While distinct cellular compartments are known to exhibit different pH environments, the observation that ∼1% of the tested RNA population adopts an i-motif conformation suggests that such structures could form *in vivo* under specific cellular conditions. Together, these findings open new avenues for exploring the structural diversity of RNA and the possible biological roles of RNA i-motifs within specific cellular contexts.

## Supporting Information

Supporting information contains UV Melting/Annealing profiles, UV difference, CD, Transitional pH plots and single molecule FRET experiments.

Source data will be deposited at DOI:10.5281/zenodo.18736033 on final publication.

## Acknowledgements

United Kingdom Biotechnology and Biological Sciences Research Council (BBSRC) [BB/W001616/1 for DG, ZW, YD and TC]; [BB/T008032/1 for MA and TC]; [BB/N022572/1 for ZW and YD]; [BB/L02229X/1 for EW and ZW] and a BBSRC White Rose DTP Studentship [2741120, for SF].

