## Supplementary data for "RNA i-Motif Formation at Neutral pH"

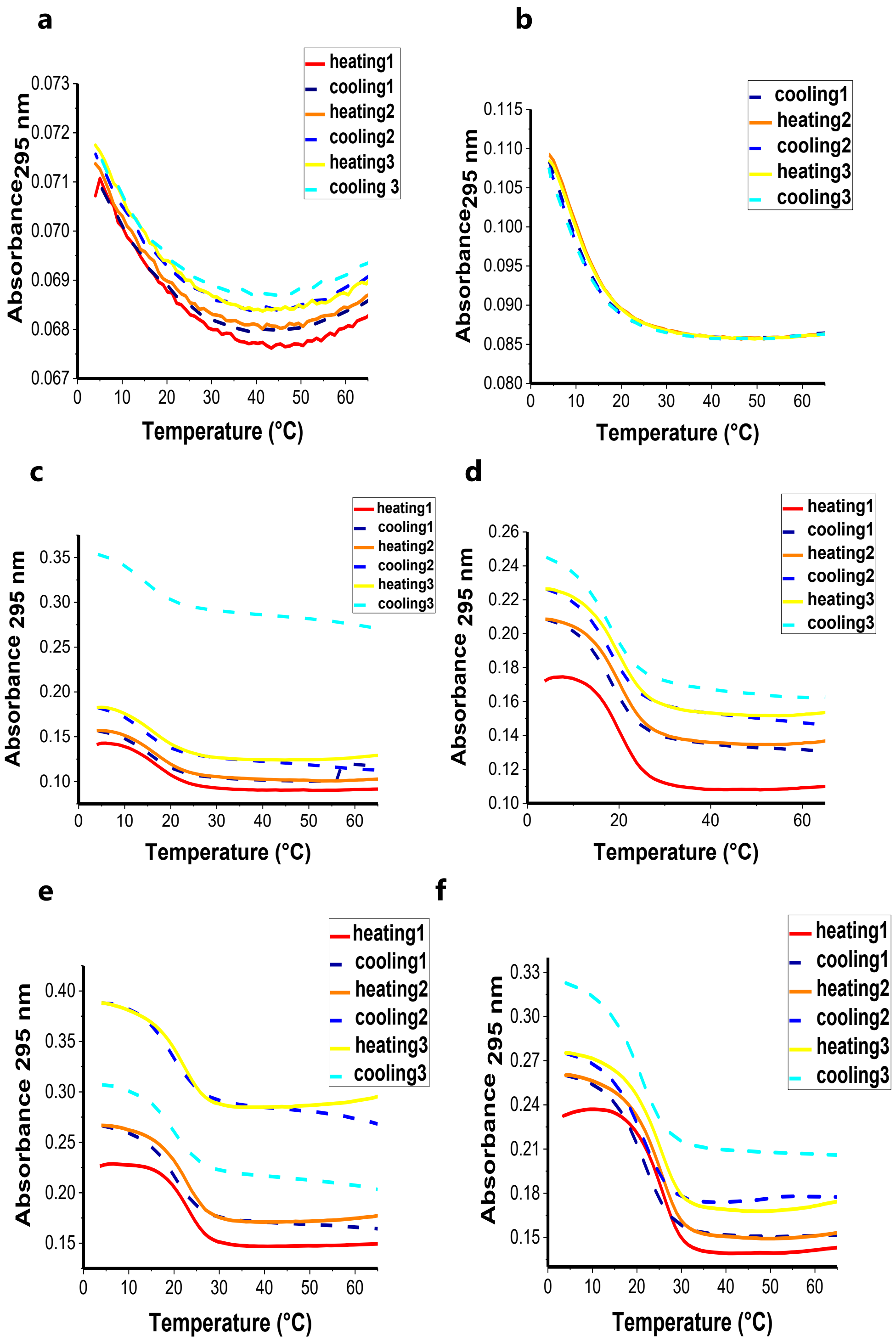

Figure S1: UV melting/annealing profiles of 2.5 μM RNA in sodium cacodylate buffer (10 mM, pH 5.5) and NaCl (100 mM) (**a**)$C_1U_3$, (**b**)$C_2U_3$, (**c**)$C_3U_3$, (**d**)$C_4U_3$, (**e**)$C_5U_3$, (**f**)$C_6U_3$.

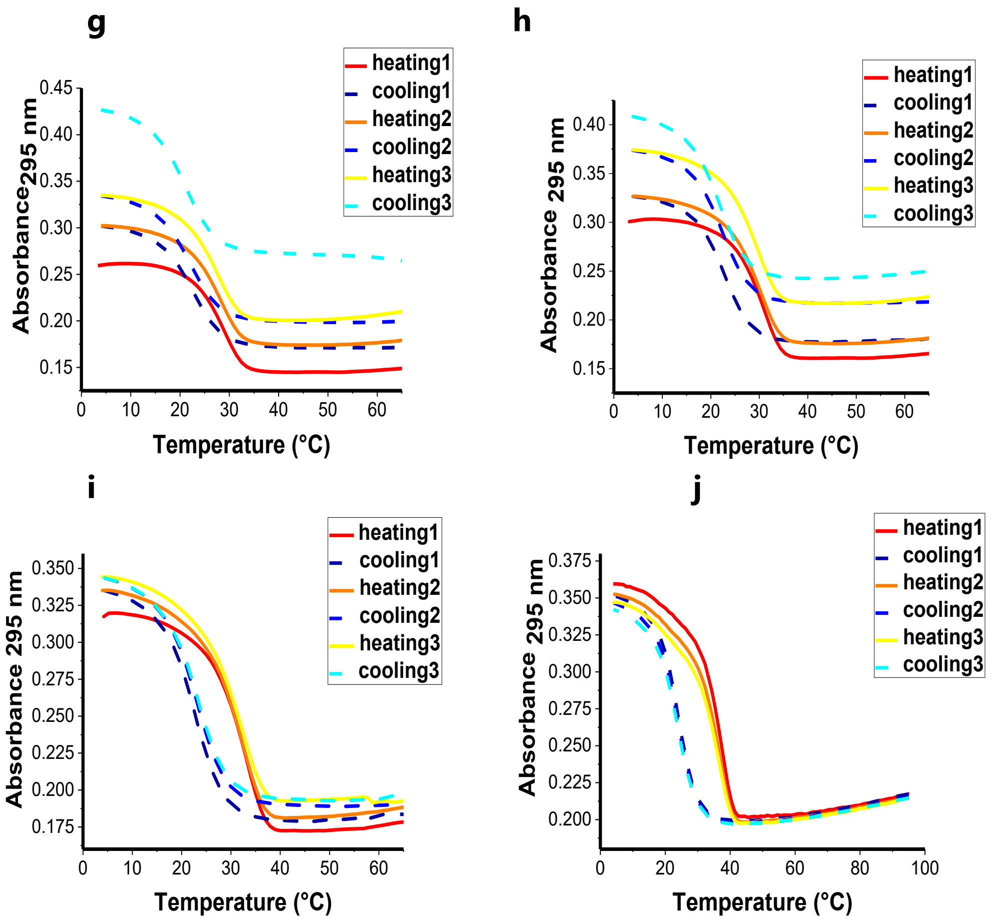

Figure S2: UV melting/annealing profiles of 2.5 µM RNA in sodium cacodylate buffer (10 mM, pH 5.5) and NaCl (100 mM) (**g**)$C_7U_3$, (**h**)$C_8U_3$, (**i**)$C_9U_3$ and (**j**)$C_{10}U_3$.

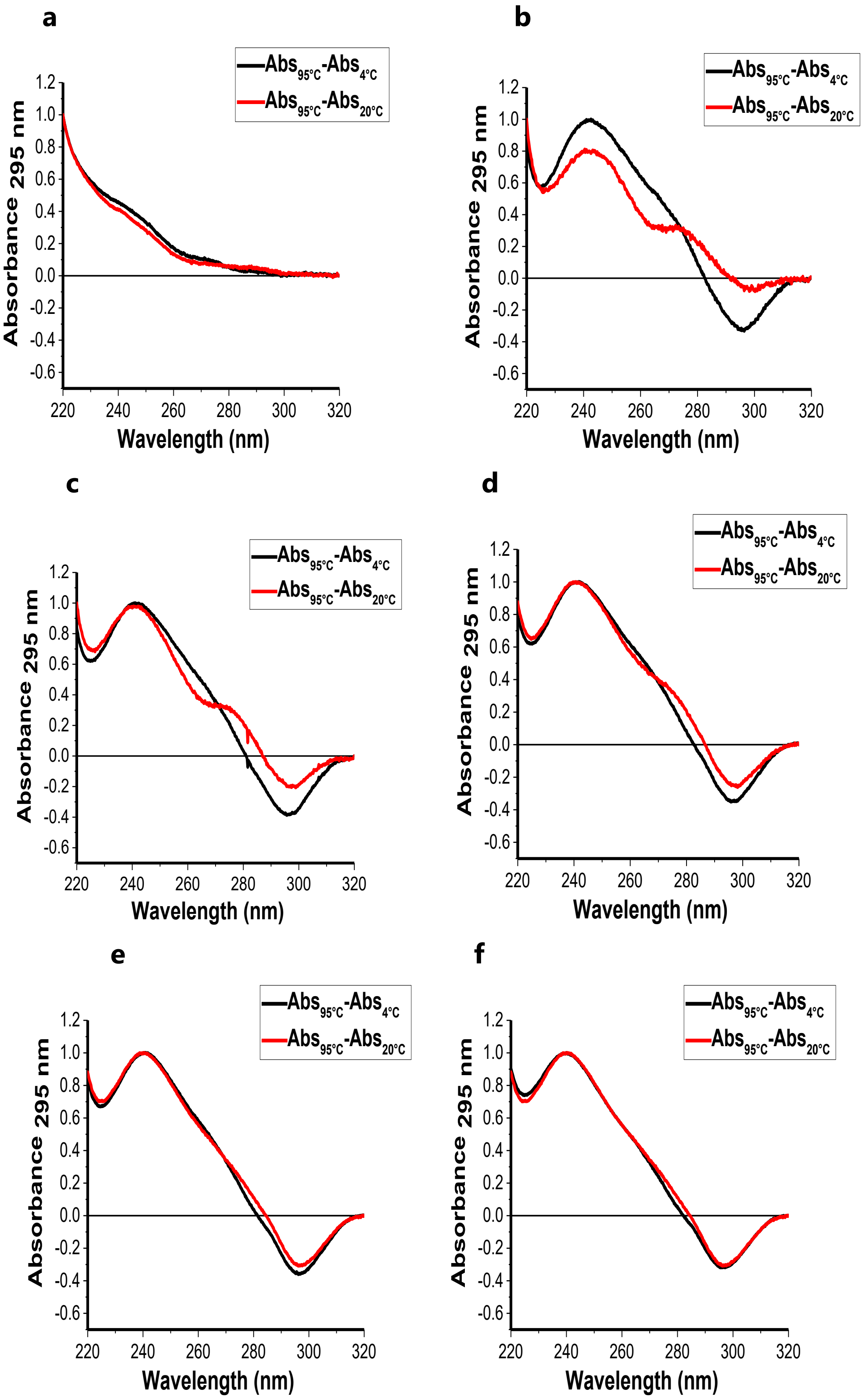

Figure S3: Thermal difference spectra profiles of 2.5 μM RNA in sodium cacodylate buffer (10 mM, pH 5.5) and NaCl (100 mM) of (**a**)$C_1U_3$, (**b**)$C_2U_3$, (**c**)$C_3U_3$, (**d**)$C_4U_3$, (**e**)$C_5U_3$, (**f**)$C_6U_3$.

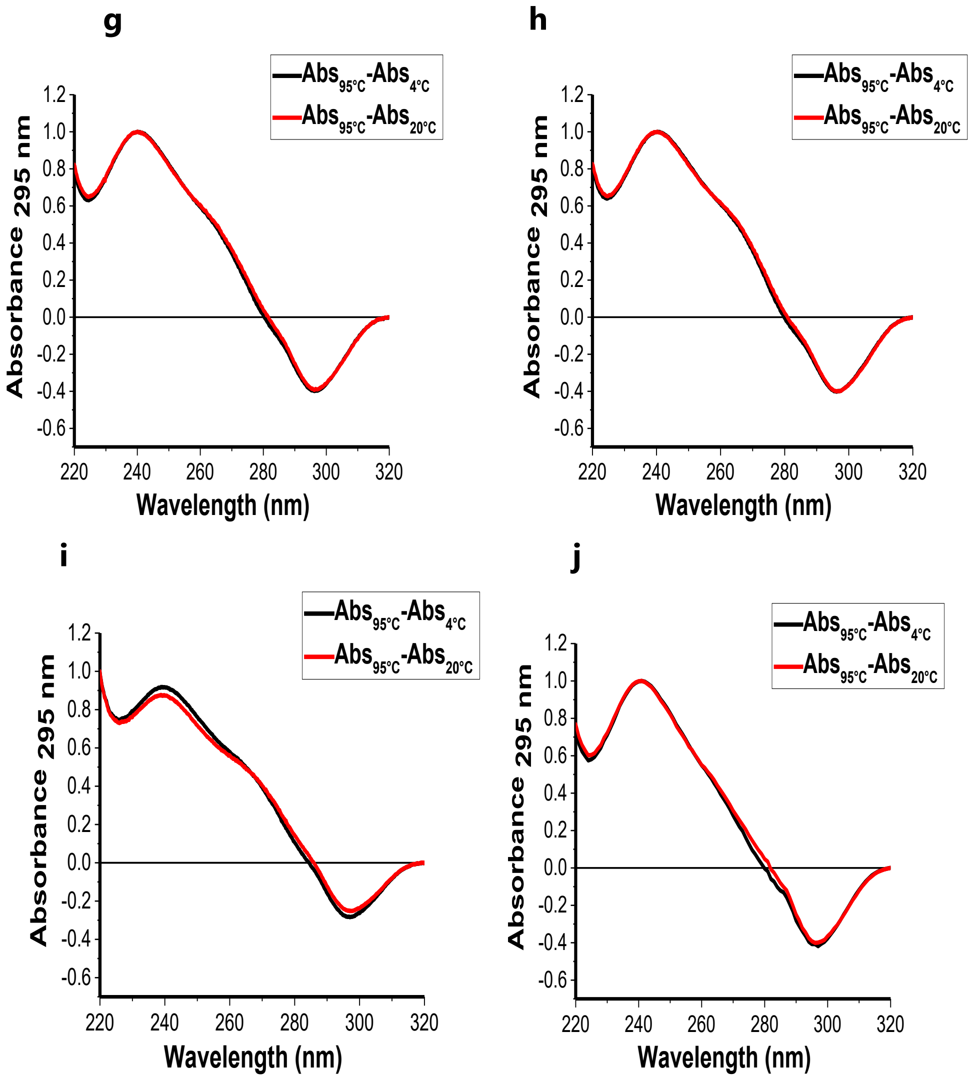

Figure S4: Thermal difference spectra (TDS) of 2.5 μM RNA in sodium cacodylate buffer (10 mM pH 5.5) and NaCl (100 mM) of (**g**)$C_7U_3$, (**h**)$C_8U_3$, (**i**)$C_9U_3$ and (**j**)$C_{10}U_3$

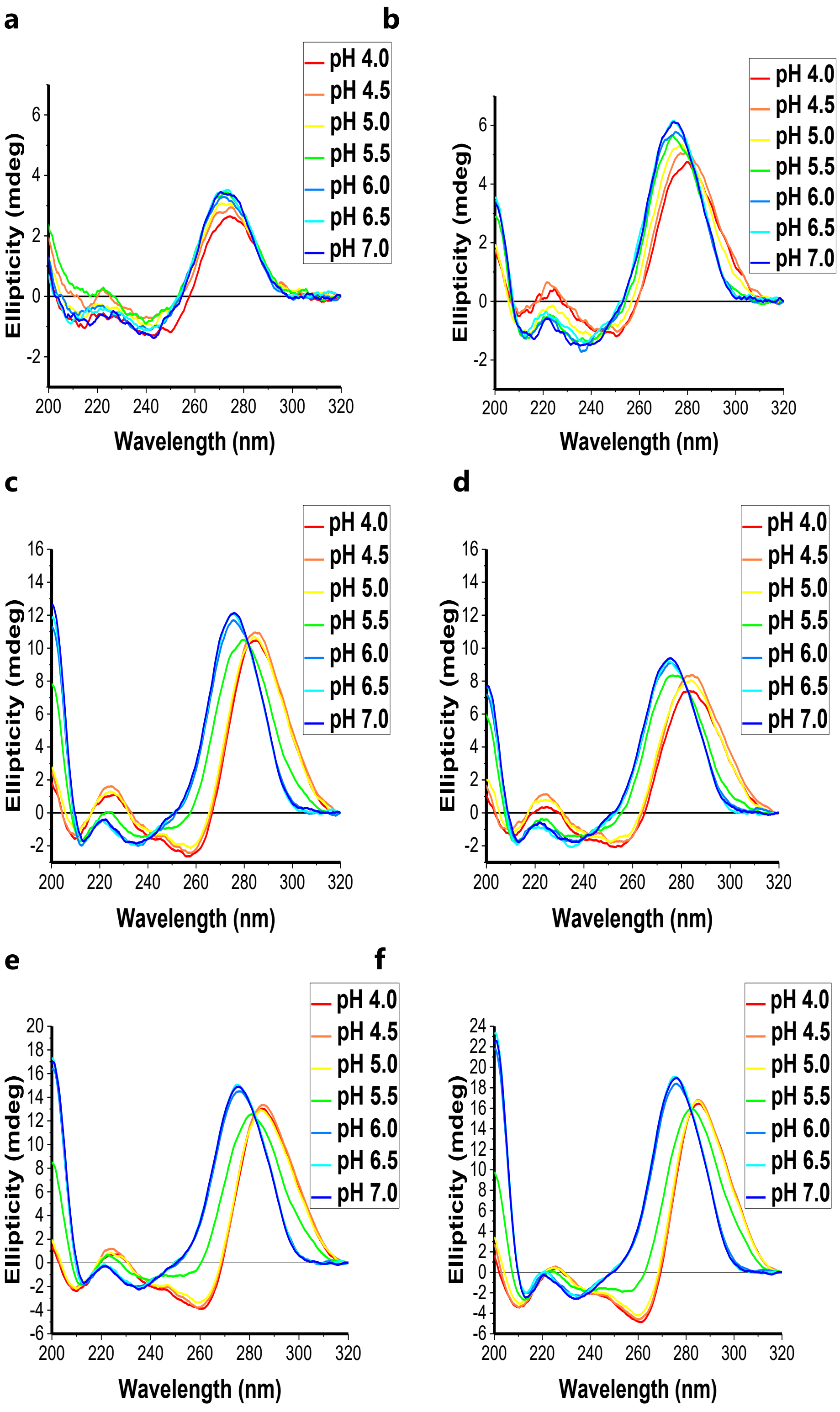

Figure S5: Circular Dichroism of 10 μM RNA in sodium cacodylate buffer (10 mM pH 5.5) and NaCl (100 mM) (**a**)$C_1U_3$, (**b**)$C_2U_3$, (**c**)$C_3U_3$, (**d**)$C_4U_3$, (**e**)$C_5U_3$, (**f**)$C_6U_3$.

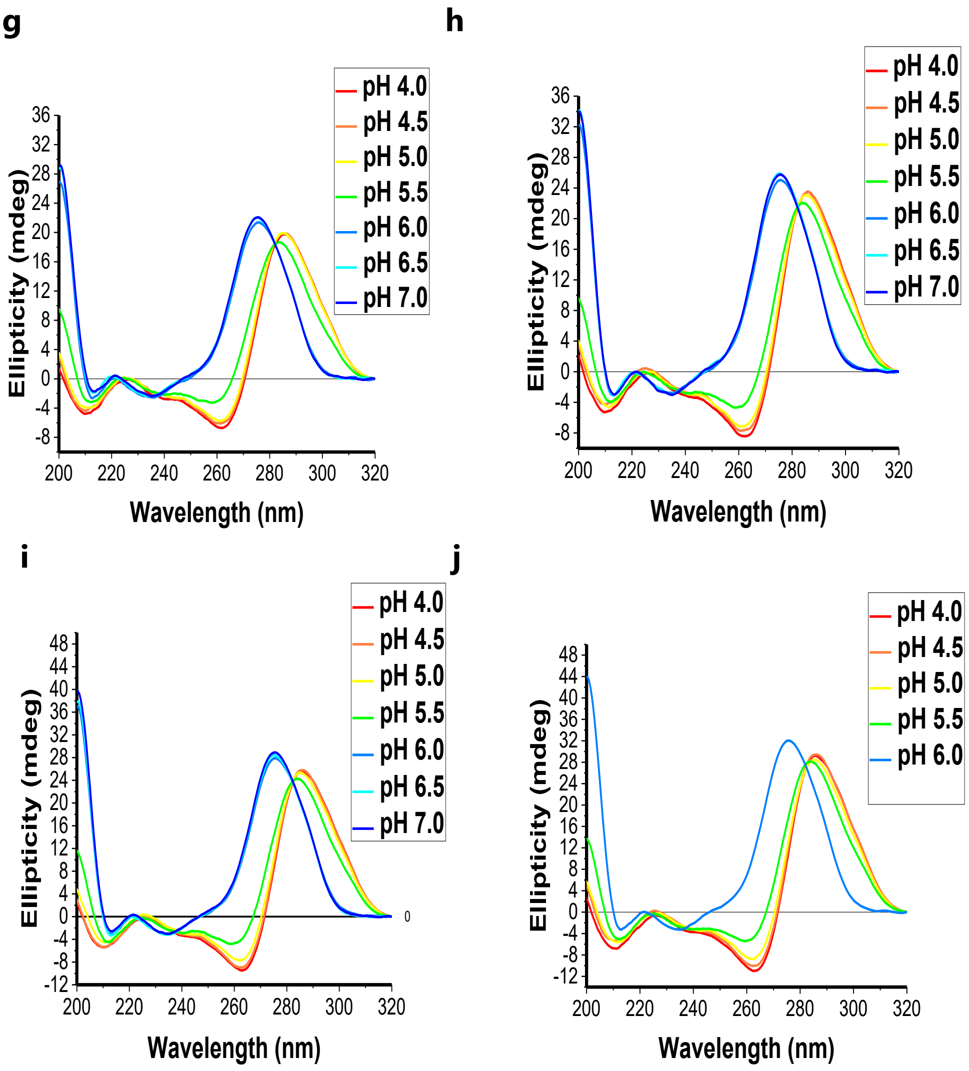

Figure S6: Circular Dichroism of 10 μM RNA in sodium cacodylate buffer (10 mM pH 5.5) and NaCl (100 mM) (**g**)$C_7U_3$, (**h**)$C_8U_3$, (**i**)$C_9U_3$, (**j**)$C_{10}U_3$.

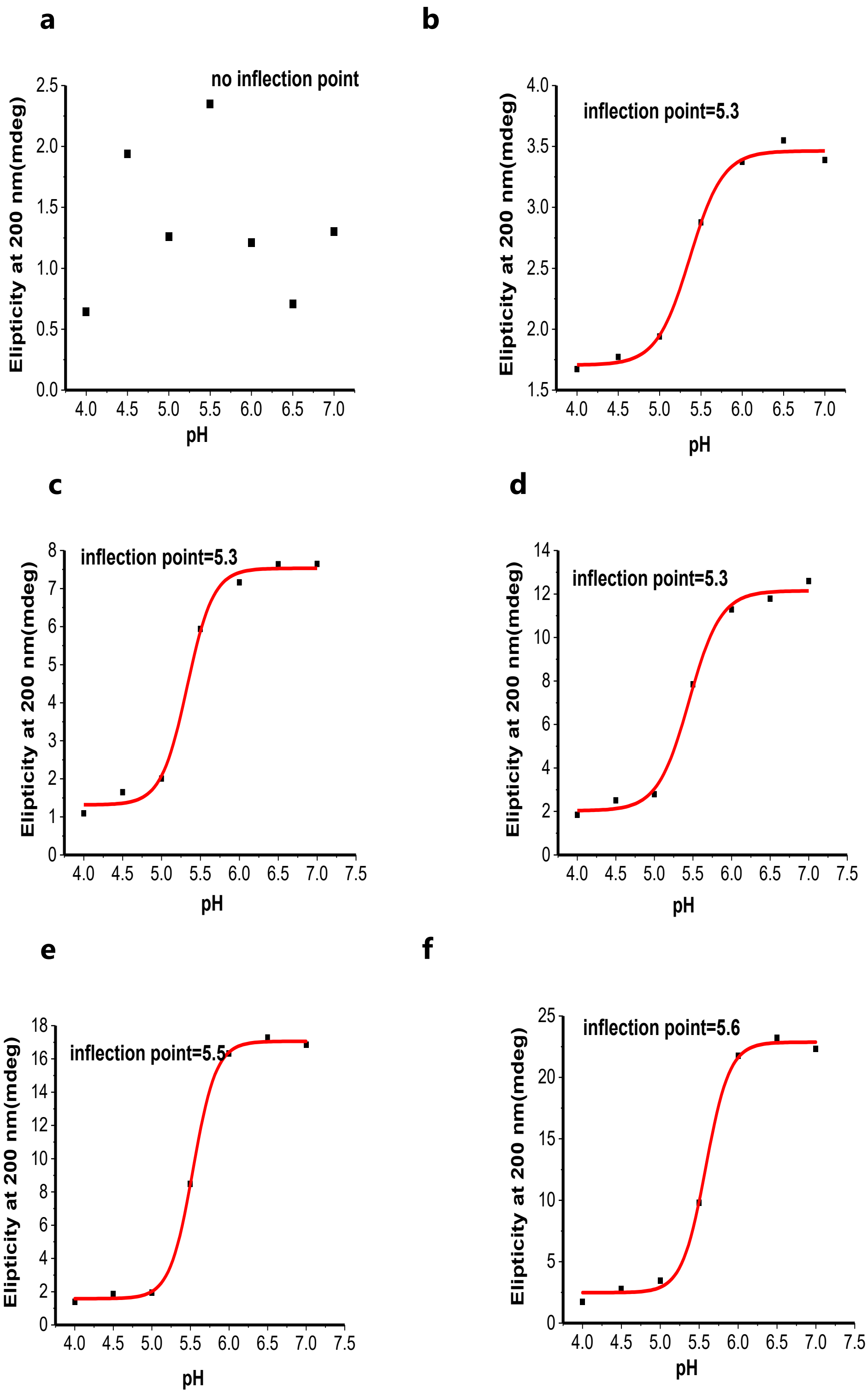

Figure S7: Ellipticity at **200** nm versus pH of 10 µM RNA in sodium cacodylate buffer (10 mM pH 5.5) and NaCl (100 mM), inflection points were calculated by plotting this data in a sigmoidal graph (**a**)$C_1U_3$, (**b**)$C_2U_3$, (**c**)$C_3$U3, (**d**)$C_4U_3$ (**e**)$C_5U_3$, (**f**)$C_6U_3$.

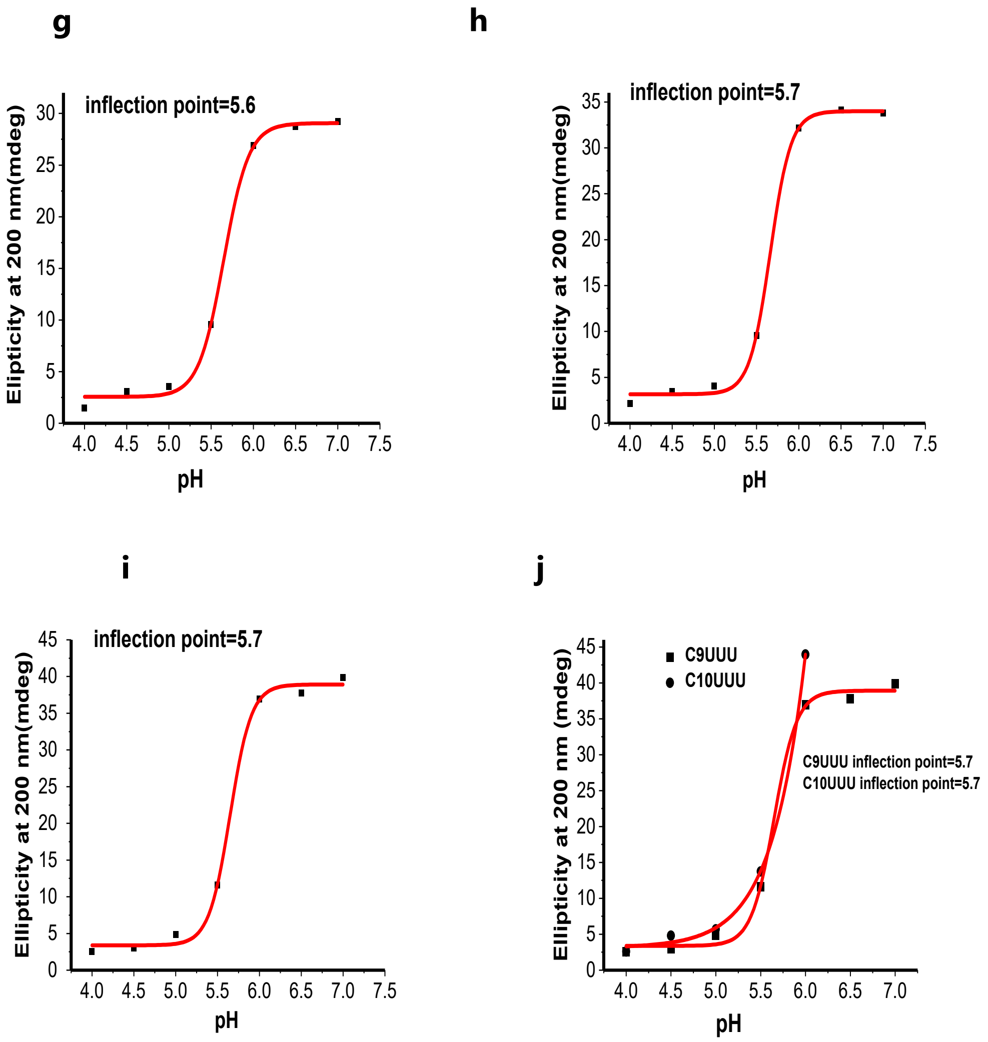

Figure S7: Ellipticity at **200** nm versus pH of 10 μM RNA in sodium cacodylate buffer (10 mM pH 5.5) and NaCl (100 mM), inflection points were calculated by plotting this data in a sigmoidal graph (**g**)$C_7U_3$, (**h**)$C_8U_3$ (**i**)$C_9U_3$, (**j**) $C_9U_3$ and $C_{10}U_3$.

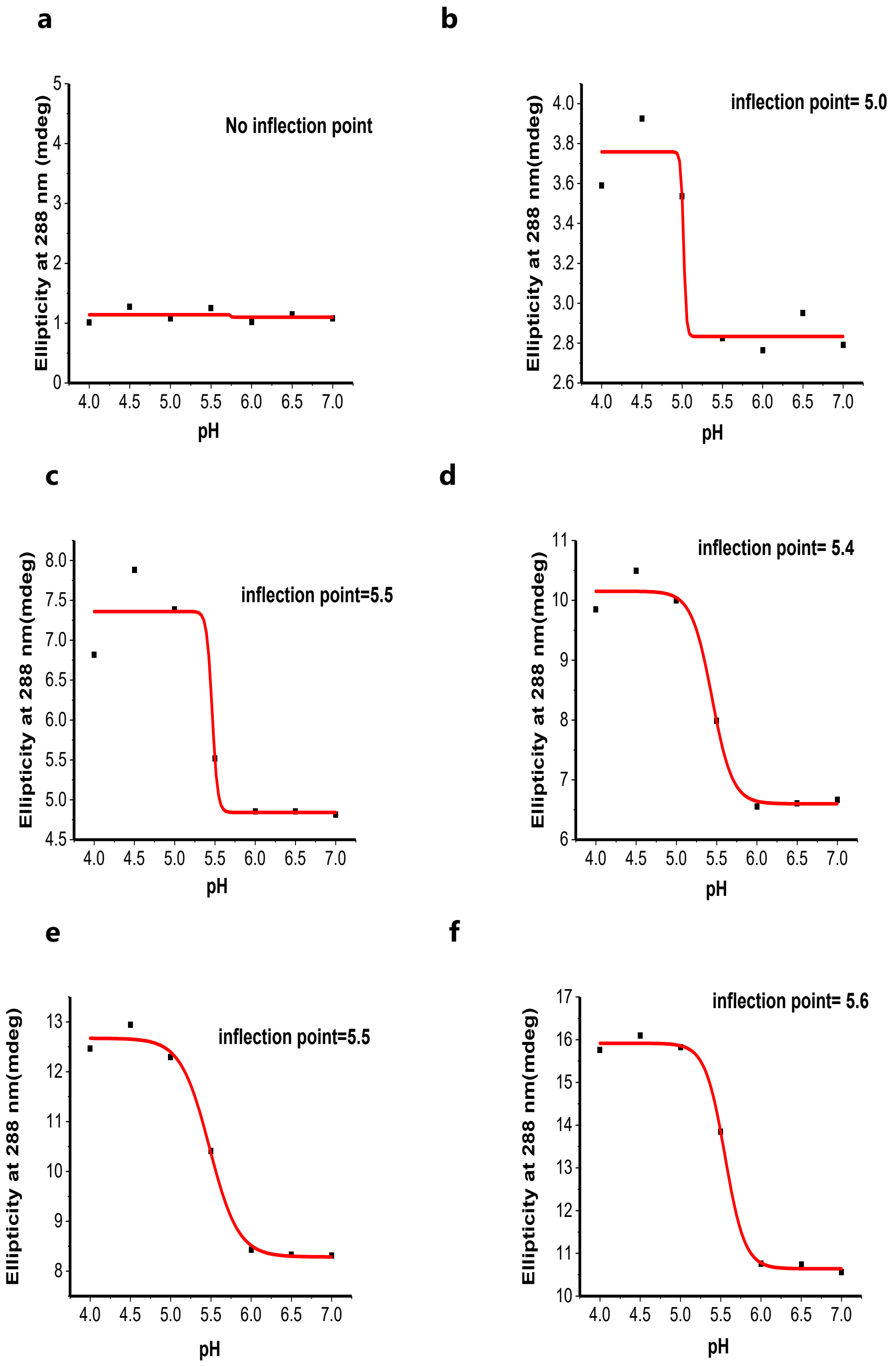

Figure S8: Ellipticity at **288** nm versus pH of 10 µM RNA in sodium cacodylate buffer (10 mM pH 5.5) and NaCl (100 mM) showing the inflection points for (**a**)$C_1U_3$, (**b**)$C_2U_3$, (**c**)$C_3$U3, (**d**)$C_4U_3$ (**e**)$C_5U_3$, (**f**)$C_6U_3$.

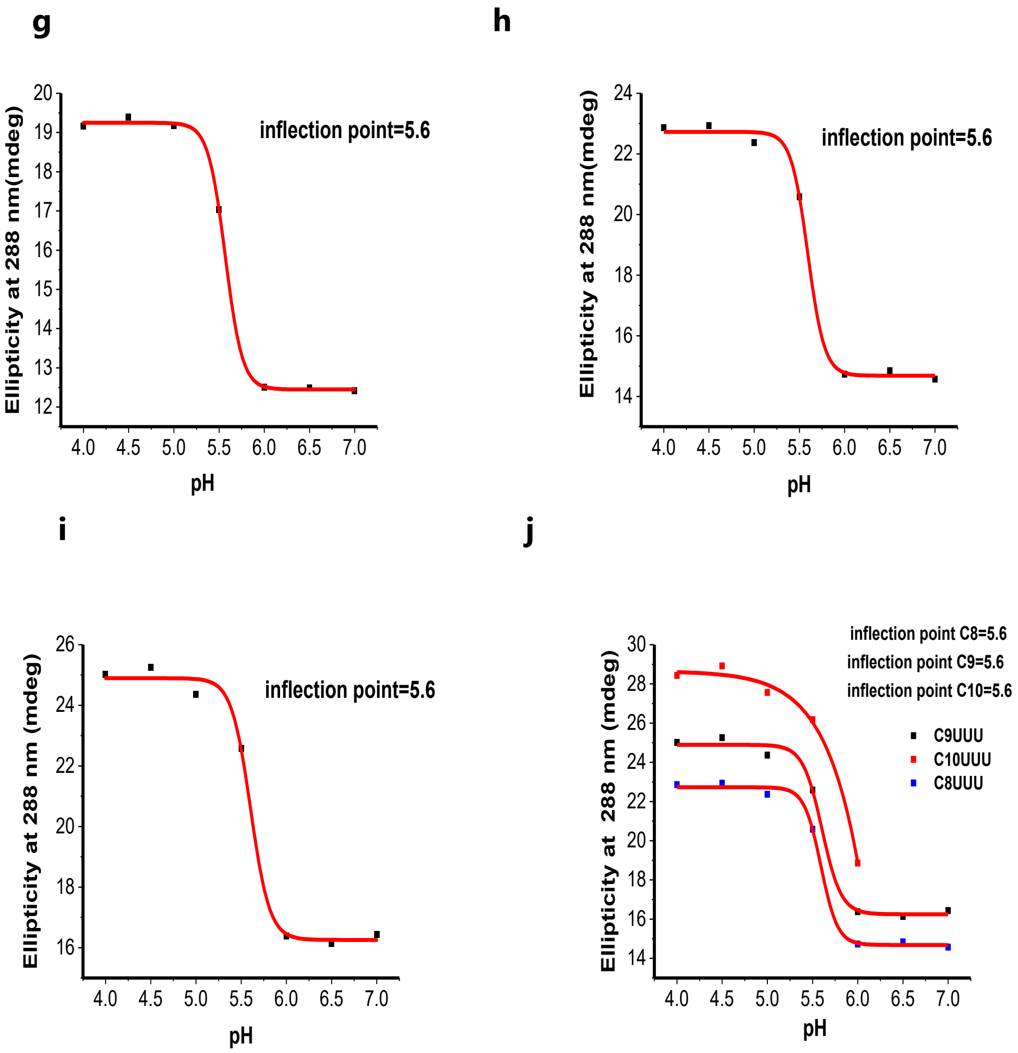

Figure S9: Ellipticity at **288** nm versus pH of 10 μM RNA in sodium cacodylate buffer (10 mM pH 5.5) and NaCl (100 mM), inflection points were calculated by plotting this data in a sigmoidal (**g**)$C_7U_3$, (**h**)$C_8U_3$, (**i**)$C_9U_3$ and (**j**)$C_8U_3$, $C_9U_3$ and $C_{10}U_3$.

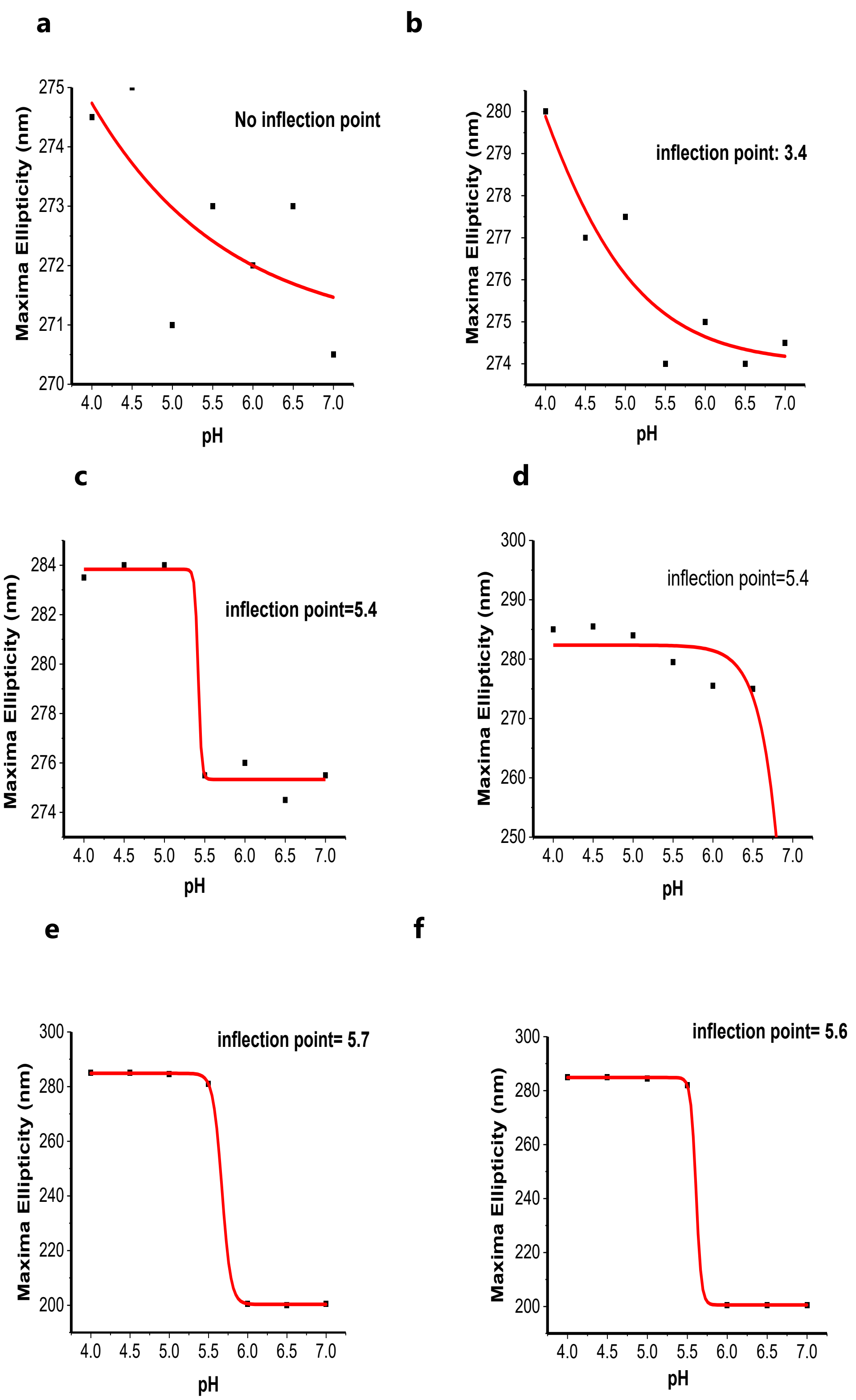

Figure S10: Plot of the wavelength of maximal ellipticity versus pH of 10 µM RNA in sodium cacodylate buffer (10 mM pH 5.5) and NaCl (100 mM), inflection points were calculated by plotting this data in a sigmoidal graph (**a**)$C_1U_3$, (**b**)$C_2U_3$, (**c**)$C_3U3$, (**d**)$C_4U_3$, (**e**)$C_5U_3$, (**f**)$C_6U_3$.

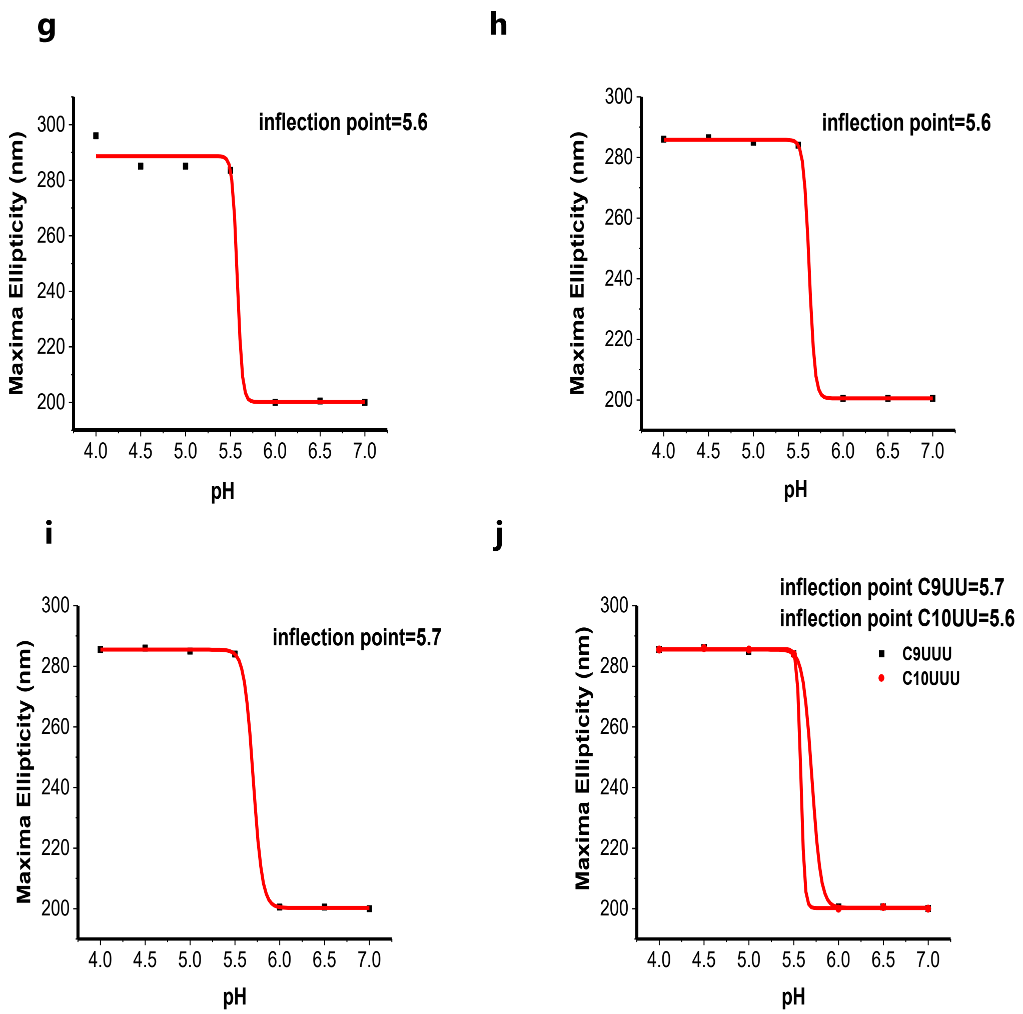

Figure S11: Plot of the wavelength of maximal ellipticity versus pH of 10 μM RNA in sodium cacodylate buffer (10 mM pH 5.5) and NaCl (100 mM), inflection points were calculated by plotting this data in a sigmoidal graph (**g**)$C_7U_3$, (**h**)$C_8U_3$, (**i**)$C_9U_3$ (**j**) $C_9U_3$ and $C_{10}U_3$.

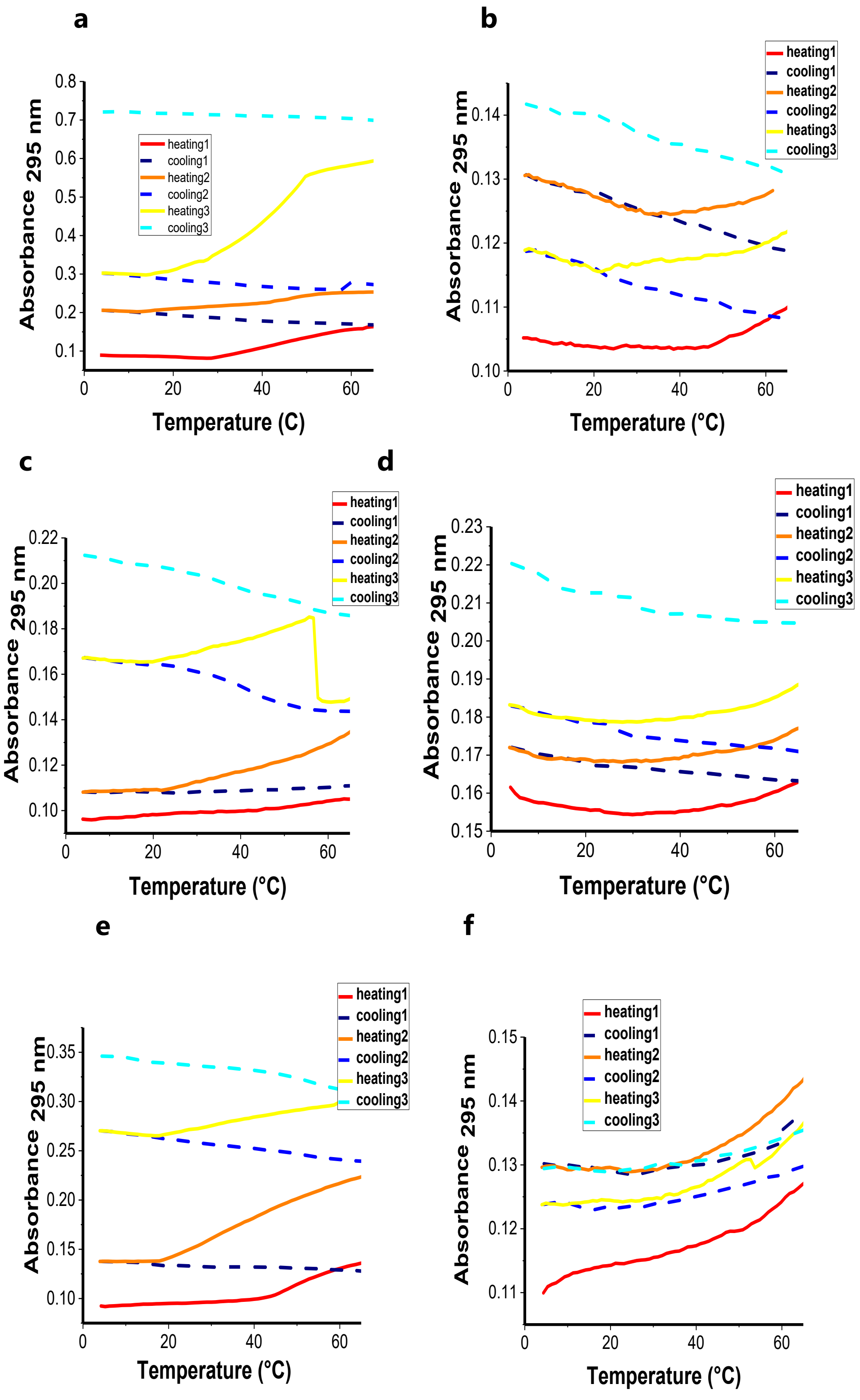

Figure S12: UV melting/annealing profiles of 2.5 μM RNA in sodium cacodylate buffer (10 mM pH 7.4) and NaCl (100 mM) (**a**)$C_1U_3$, (**b**)$C_2U_3$, (**c**)$C_3U_3$, (**d**)$C_4U_3$, (**e**)$C_5U_3$, (**f**)$C_6U_3$.

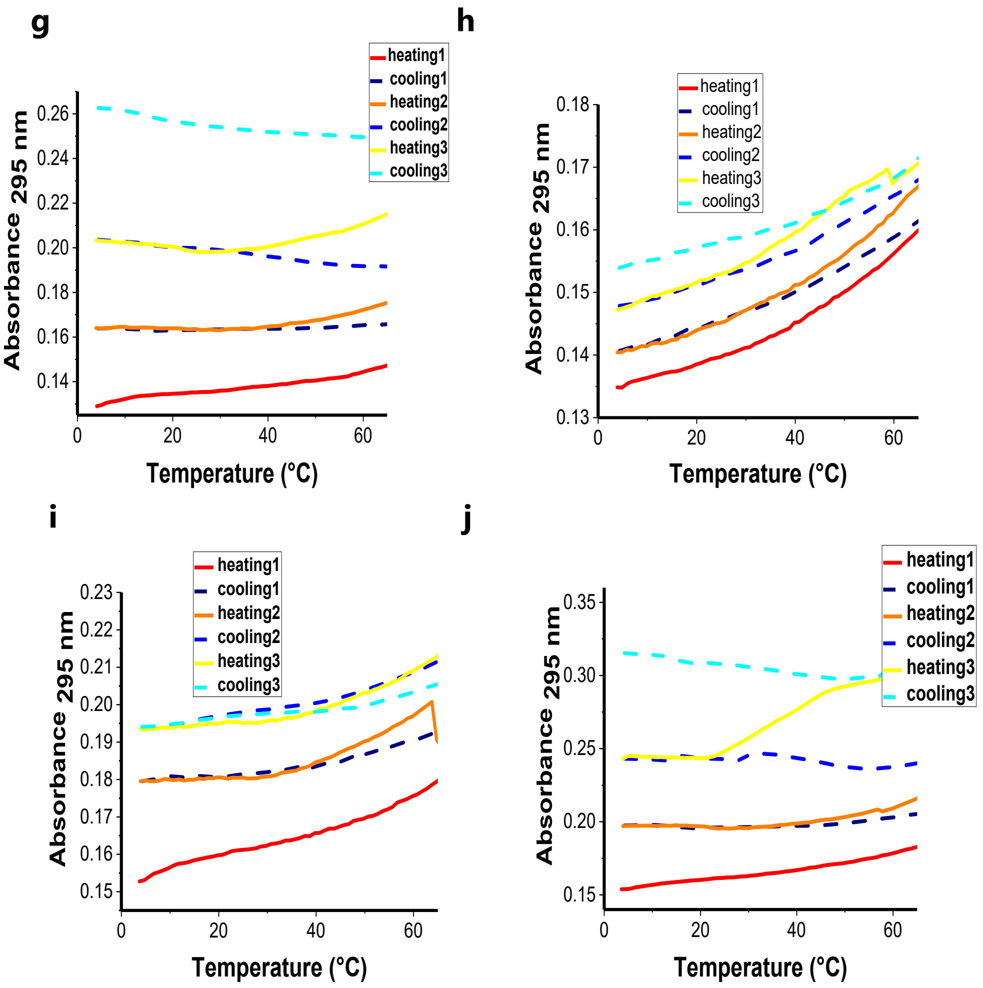

Figure S13: UV melting/annealing profiles of 2.5µM RNA in sodium cacodylate buffer (10 mM pH 7.4) and NaCl (100 mM) (**g**)$C_7U_3$, (**h**)$C_8U_3$, (**i**)$C_9U_3$ and (**j**)$C_{10}U_3$.

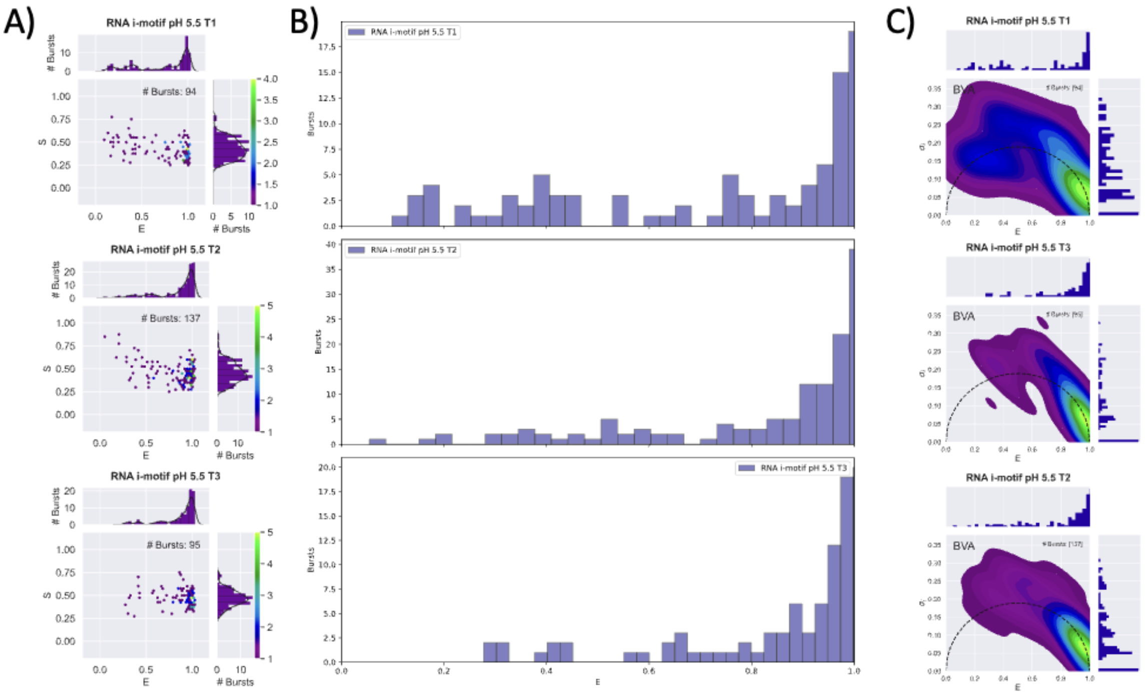

Figure S14: Single molecule FRET measurement with C5U3$_{FRET}$ at pH 5.5 for 1 hour in triplicate. A) 2D-Histogram of dual channel burst search of each of the repeats B) Histogram of FRET Efficiency from the dual channel burst search C) Burst variance analysis (BVA) a method that detects within-burst dynamics by comparing the standard deviation of the FRET efficiency of bursts to expected – higher standard deviation indicates the burst giving that FRET efficiency is dynamic (i.e. moving) while traversing the confocal volume; lower (below dashed line) indicates it is static (i.e. likely in some "fixed" state/conformation)

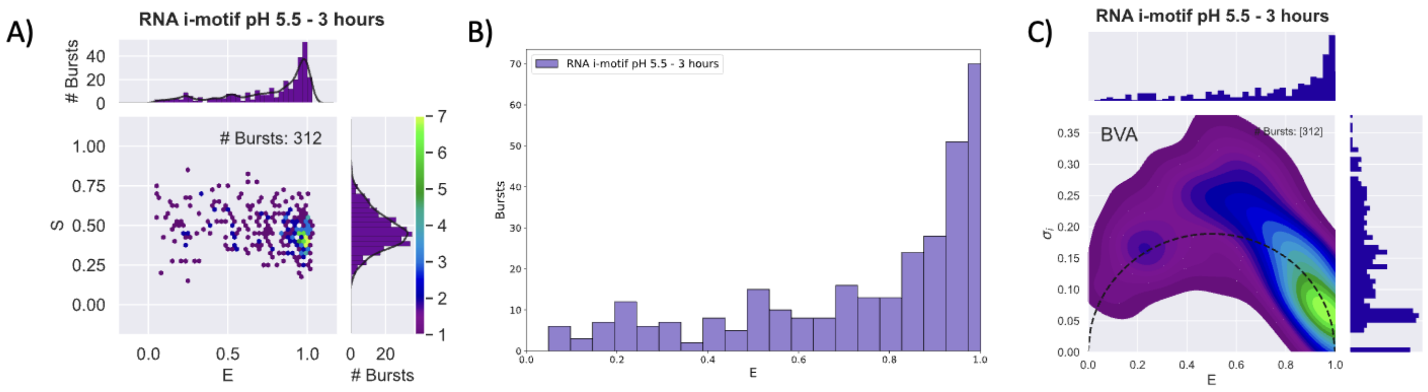

Figure S15: Single molecule FRET measurement with $C5U3_{FRET}$ at pH 5.5 for 3 hours. A) 2D-Histogram of dual channel burst search B) Histogram of FRET Efficiency from the dual channel burst search C) Burst variance analysis (BVA) a method that detects within-burst dynamics by comparing the standard deviation of the FRET efficiency of bursts to expected – higher standard deviation indicates the burst giving that FRET efficiency is dynamic (i.e. moving) while traversing the confocal volume; lower (below dashed line) indicates it is static (i.e. likely in some ″fixed″ state/conformation)

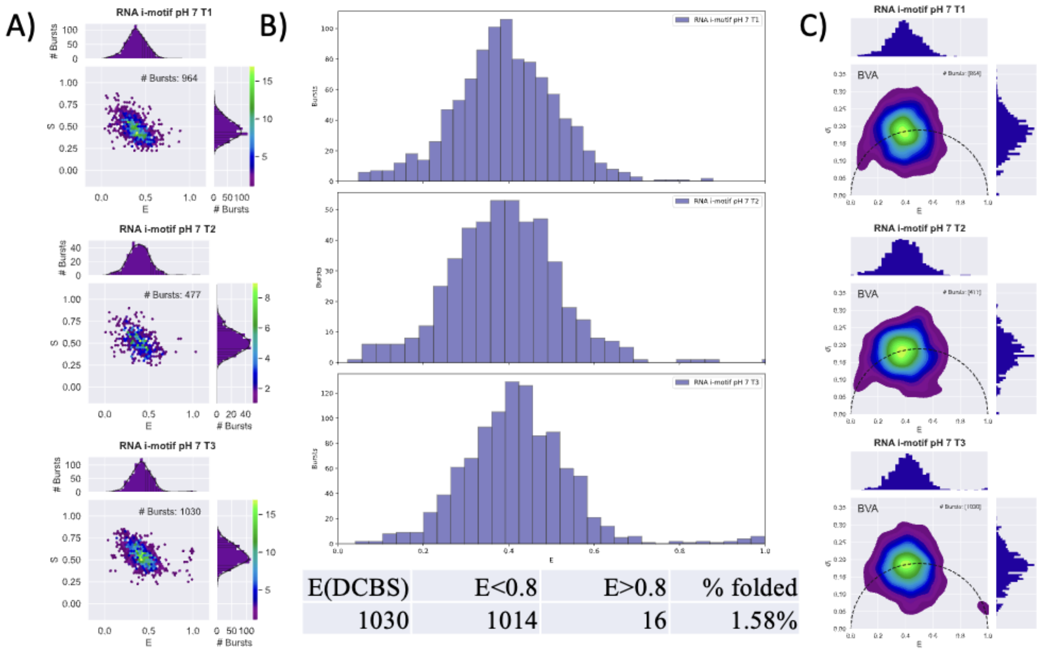

| E(DCBS) | E<0.8 | E>0.8 | % folded |
|---|---|---|---|
| 1030 | 1014 | 16 | 1.58% |

Figure S16: Single molecule FRET measurement with $C5U3_{FRET}$ at pH 7 for 1 hour in triplicate. A) 2D-Histogram of dual channel burst search of each of the repeats B) Histogram of FRET Efficiency from the dual channel burst search C) Burst variance analysis (BVA) a method that detects within-burst dynamics by comparing the standard deviation of the FRET efficiency of bursts to expected – higher standard deviation indicates the burst giving that FRET efficiency is dynamic (i.e. moving) while traversing the confocal volume; lower (below dashed line) indicates it is static (i.e. likely in some "fixed" state/conformation)

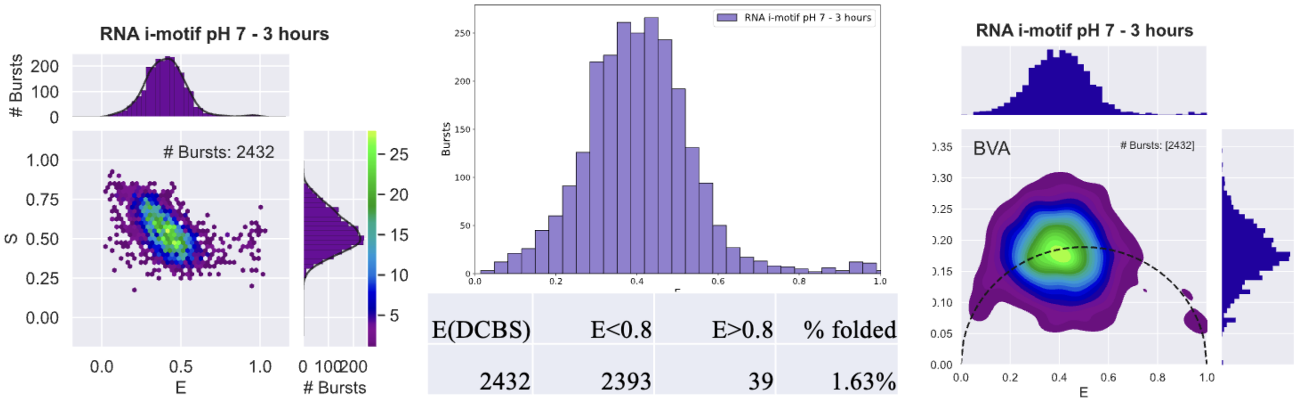

| E(DCBS) | E<0.8 | E>0.8 | % folded |
|---|---|---|---|
| 2432 | 2393 | 39 | 1.63% |

Figure S17: Single molecule FRET measurement with $C5U3_{FRET}$ at pH 7 for 3 hours. A) 2D-Histogram of dual channel burst search B) Histogram of FRET Efficiency from the dual channel burst search C) Burst variance analysis (BVA) a method that detects within-burst dynamics by comparing the standard deviation of the FRET efficiency of bursts to expected – higher standard deviation indicates the burst giving that FRET efficiency is dynamic (i.e. moving) while traversing the confocal volume; lower (below dashed line) indicates it is static (i.e. likely in some "fixed" state/conformation)

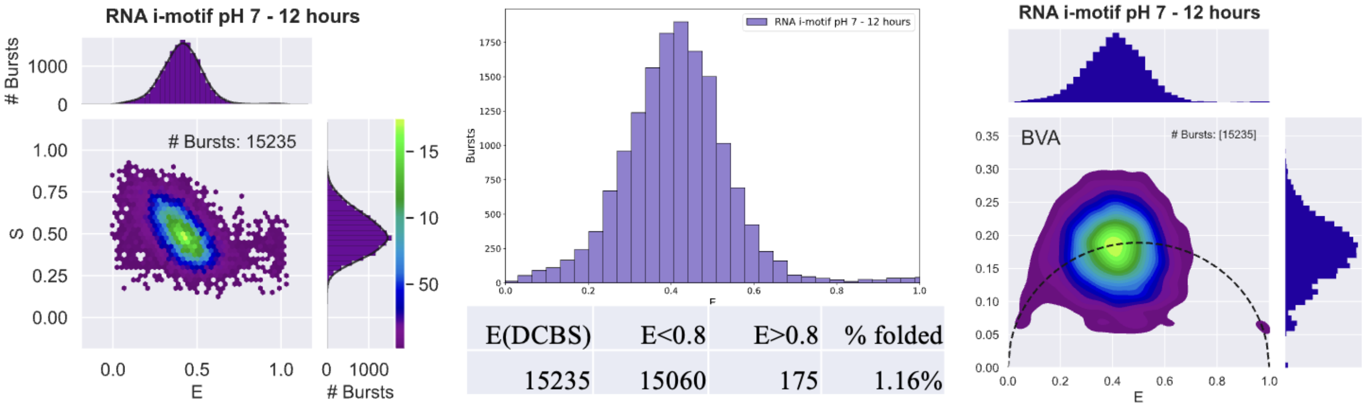

| E(DCBS) | E<0.8 | E>0.8 | % folded |
|---|---|---|---|
| 15235 | 15060 | 175 | 1.16% |

Figure S18: Single molecule FRET measurement with C5U3$_{FRET}$ at pH 7 for 12 hours. A) 2D-Histogram of dual channel burst search B) Histogram of FRET Efficiency from the dual channel burst search C) Burst variance analysis (BVA) a method that detects within-burst dynamics by comparing the standard deviation of the FRET efficiency of bursts to expected – higher standard deviation indicates the burst giving that FRET efficiency is dynamic (i.e. moving) while traversing the confocal volume; lower (below dashed line) indicates it is static (i.e. likely in some "fixed" state/conformation)
